# STF-62247 blocks late stages of autophagy by disrupting lysosomal physiology

**DOI:** 10.1101/298729

**Authors:** Nadia Bouhamdani, Dominique Comeau, Kevin Cormier, Sandra Turcotte

## Abstract

STF-62247 was previously identified as a promising compound able to selectively target the loss of the tumor suppressor gene von Hippel-Lindau (VHL) in renal cell carcinomas. This present work investigates the effect of STF-62247 on the autophagic flux. Our investigations show that STF-62247 blocks late stages of autophagy through lysosomal disruption. Indeed, STF-62247 localizes at lysosomes and causes unregulated swelling of these acidic compartments in VHL-mutated cells, linking a potential role for VHL in lysosomal integrity. Knock-outs of BECN1 and ATG5 were able to rescue the viability of VHL-mutated cells in response to STF-62247 but did not rescue the lysosomal swelling. In fact, neutralizing the lysosomal pH by inhibiting the vacuolar H+-ATPase completely rescued this phenotype. Moreover, we show that STF-62247 disrupts endocytic routes and causes cathepsin D trafficking defects. This mechanistic study is the first to characterize STF-62447 as a novel lysosomotropic compound. Importantly, our study re-classifies STF-62247 as a blocker of later stages of autophagy and highlights its potential usage as a powerful new tool for endocytic and autophagy-related research.

## INTRODUCTION

Renal cell carcinoma (RCC) is the most common type of kidney cancer and is considered the most lethal of all genitourinary cancers. The majority of RCC diagnoses are of clear cell subtype (ccRCC) and accounts for 80-90% of renal epithelial malignancies.(1) Alarmingly, up to 20% of patients present with locally advanced or metastatic disease and more than a third of patients with early-stage disease treated with curative intent will face metastatic recurrence.(2, 3) Clear cell renal tumors are frequently associated with a loss of function of the von Hippel-Lindau gene (VHL) which leads to stabilization of hypoxia inducible factors alpha (HIF-*α*).(4) Thus, upon VHL loss, upregulation of proangiogenic genes, uncontrolled cell growth and pH deregulation make ccRCCs highly vascularized and aggressive. There are several targeted therapies available for the treatment of metastatic RCC such as tyrosine kinase and mammalian target of rapamycin (mTOR) inhibitors as well as anti-vascular epithelial growth factor antibodies.(5–7) Unfortunately, development of tumor resistance renders their clinical response short-lived and consequently, metastatic RCC remains today an unmet clinical challenge.

Macroautophagy (hereafter referred to as autophagy) is a highly conserved catabolic process that assures cellular homeostasis. Canonical autophagy is characterized by the formation of a phagophore membrane that elongates and closes to form a double-membrane vesicle called autophagosome (AP). The enclosed intracellular components are delivered to lysosomes for degradation. This process can be dissected in four steps; *1)* initiation; *2)* nucleation; *3)* maturation and *4)* fusion. The initiation and nucleation phases are essential for phagophore formation and are regulated by the complex ULK1-mATG13-FIP200 and by a complex including the class 3 phosphatidylinositol-3-kinase (VPS34), p150, ATG14L and Beclin-1, respectively. The maturation phase, consisting in phagophore elongation, curvature and closure is regulated by ubiquitin-like conjugation systems ensuring the lipidation of the AP marker LC3 (LC3-II) and its insertion in the membrane via the complex ATG12-ATG5-ATG16L1. The adaptor protein p62/sequestosome-1(SQSTM1) binds LC3-II and facilitates autophagic degradation of ubiquitinated cargo. Its increased levels often correlate with autophagy inhibition.

In cancer, autophagy can serve as a survival pathway, especially in advanced stages. It is thought to promote tumor growth and progression by enhancing cancer cells’ survival in harsh conditions such as hypoxia and nutrient limitations.(8) In this context, suppressing autophagy has shown promise as a therapeutic modality to sensitize cancer cells to conventional and targeted therapies.(9) However, given the multifarious behaviors of each cancer, fine-tuning specific mechanisms of autophagic suppression remains difficult. Indeed, autophagy and its interconnected pathways seem to be uniquely regulated in each cancer. Thus, novel autophagy-modulating tools are needed to unravel unique mechanistic insights existing between autophagy, endocytosis and vesicular trafficking pathways. One of the first studies to suggest autophagy-modulation as a potential therapy for VHL-deficient RCCs did so by identifying STF-62247, a compound demonstrating selectivity for RCC cells harboring VHL mutations.(10) Considering that the extensive study of autophagy was not the primary goal of this previous paper, this present work aimed at an in-depth study of STF-62247’s effects on the autophagic flux. We show that STF-62247 is a blocker of late stages of autophagy by targeting lysosomal physiology. We highlight STF-62247’s lysosomotropic properties causing enlargement of endo-lysosomal compartments and vesicle trafficking defects. Finally, our study elucidates a potential new role for VHL in maintaining lysosomal integrity and highlights its potential usage as a powerful new tool for the study of vesicle trafficking and lysosome-related research.

## RESULTS

### STF-62247 blocks late stages of autophagy independently of VHL status

Three cell line models have been chosen for this study; VHL-mutated ccRCC cell line (RCC4), its subclone counterpart stably transfected with an expression vector encoding VHL (RCC4 VHL) and a cell line containing endogenous VHL in which canonical autophagy has been extensively studied (HeLa).(11–14) To assess the selective properties of STF-62247 (STF), the sensitivity of all three cell lines has been tested by XTT viability assays (**Fig.1A)**. VHL-mutated RCC4 cells are significantly more sensitive to the small compound compared to VHL-containing cell lines RCC4 VHL and HeLa. It should be noted that HeLa cells remain relatively unaffected when compared to RCC4 VHL cells, even at high STF concentrations such as 10 and 20*μ*M (**Fig.1A)**. In addition to the selective properties of STF, a striking phenotype is observed following treatment; the formation of large translucid intracytoplasmic vacuoles (**Fig. 1B**). The vacuolar phenotype of VHL-containing cell models (RCC4VHL, HeLa) remains controlled with modest vacuole sizes whereas VHL-defective RCC4 cells present continuously expanding vacuoles (**Fig. 1B**).

**Figure 1.**
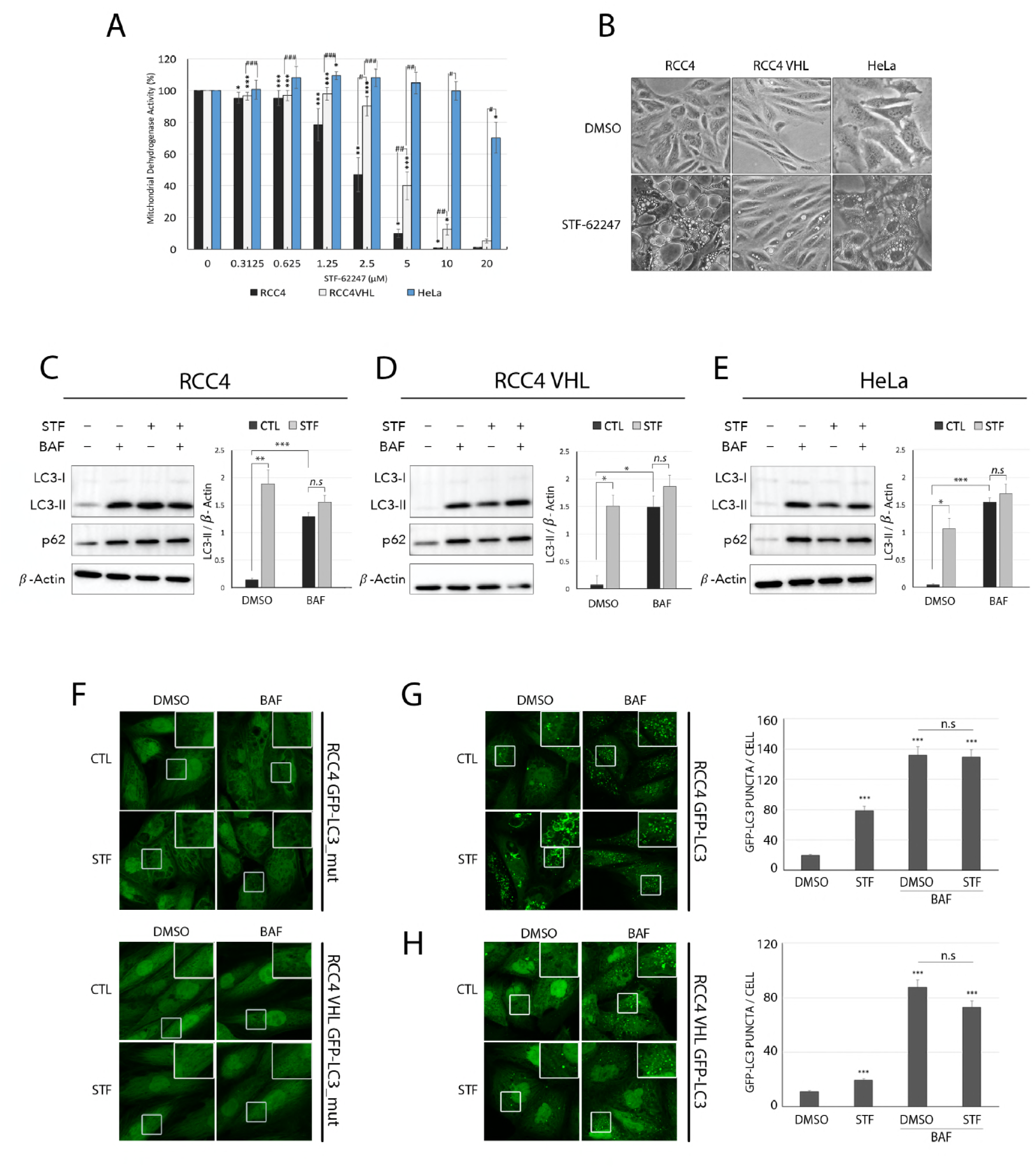
STF-62247 is a blocker of late stages of autophagy independently of VHL status. (A) XTT viability assay of RCC4, RCC4 VHL and HeLa cell models in response to 0-20*μ*M of STF. **(B)** Inverted-light microscopy images of vacuolization phenotype of RCC4, RCC4 VHL and HeLa cell models in response to 48 hr of STF treatment. Immunoblot analysis with quantification following 4 hr of STF and BAF treatment alone, or combined in **(C)** RCC4, **(D)** RCC4 VHL and **(E)** Hela. **(F)** RCC4 and RCC4VHL stably expressing a lipidation-defective GFP-LC3 (glycine 120 changed to an alanine residue) control and treated as in (C-E). Confocal microscopy of stably expressing GFP-LC3 with puncta quantification of **(G)** RCC4 and **(H)** RCC4 VHL and treated as in (C-E). Mean and SEM were calculated from at least three independent experiment. Statistical analysis compared untreated cells with treated cells (*p<0.05, **p<0.01, ***p<0.001) and the difference between the two cell lines (#p<0.05, ##p<0.01, ###p<0.001) at each concentration using Student’s T-test.

In order to evaluate STF-62247’s effect on autophagy and to identify possible related differences that would correlate with VHL-dependent viability, LC3-turnover assays were performed (**Fig. 1C-1E**). Levels of LC3-II and p62 were measured by immunoblot analysis in response to STF and to a saturating concentration of the lysosomal-disrupting agent Bafilomycin A1 (BAF) either alone, or in combination with STF. BAF, a known inhibitor of vacuolar H+ ATPase pump (V-ATPase) is used as a control for its known inhibitory effects on autophagy efflux.(15–17) BAF control alone resulted in significant accumulation of LC3-II and p62 levels in every cell line model, as expected (**Fig.1C-1E**). The combination of BAF plus STF did not show an additive effect on LC3-II levels compared to BAF alone in neither VHL-mutated RCC4 cells nor in cell lines containing a functioning gene, RCC4 VHL and HeLa (**Fig. 1C-1E**). To confirm these results, cell lines expressing stable GFP-LC3 were generated (RCC4.GFP-LC3 and RCC4VHL.GFP-LC3) and puncta formation by confocal microscopy was quantified (**Fig.1F-1H**). Importantly, a lipidation-defective GFP-LC3 mutant where glycine 120 is mutated to alanine (RCC4 GFP-LC3_mut and RCC4VHL GFP-LC3_mut) was utilized as a specificity control in order to avoid possible aggregation of the chimera, unrelated to autophagy that would be due to high levels of expression **(Fig. 1F)**.(18, 19) In response to BAF or STF, GFP-LC3-mut models show no puncta formation confirming the functionality of the chimeric models (**Fig. 1F**). In agreement with immunoblot analyses, combination treatments of BAF plus STF did not result in an additional increase in GFP-LC3 puncta compared to BAF treatment alone in both RCC4.GFP-LC3 and RCC4VHL.GFP-LC3 models (**Fig. 1G-1H**). Together, these observations reveal STF-62247 to be a blocker of autophagy rather than an inducer, as was previously described.(10, 20, 21) Furthermore, these results strongly suggest that this compound blocks terminal stages of autophagy independently of VHL status.

### Long-term STF-62247 treatment decreases the number of endo-lysosomal vesicles in VHL-mutated RCC4 cells

To further evaluate the effects of STF-62247 on the autophagic flux, a tandem monomeric mCherry-GFP-tagged LC3 was stably expressed in RCC4, RCC4VHL and HeLa cell lines (**Fig.2A-2C**). Quickly, yellow fluorescence is associated with autophagic vesicles (AV) while red fluorescence alone decorates lower-pH organelles such as autolysosomes (AL).(22, 23) In accordance with the above-mentioned GFP-LC3 puncta count, combinatory treatment of BAF plus STF did not show any additive effects in the numbers of autophagic vesicles (**Fig 1C-1E and 2A-2C**). Interestingly, STF treatment alone increased the numbers of autophagic vesicles in all three cell lines but not those of autolysosomes, confirming a block in the flux (**Fig.2A-2C**). In fact, the number of autolysosomes remained relatively unchanged in VHL-mutated RCC4 cells whereas a significant decrease in autolysosomes was observed in VHL-proficient cell lines (**Fig.2B, 2C**). To be rigorous, levels of endogenous LC3 and LAMP-1 (marker of endo-lysosomal structures) were also quantified by immunofluorescence following a prolonged STF treatment (**Fig.2D, 2E**). In response to 72 hr of STF treatment, VHL-defective RCC4 cells do not restore their levels of endo-lysosomal structures and accumulate APs (**Fig. 2D, 2E**). Contrastingly and in accordance with VHL-proficient RCC cells’ capacity to overcome the vacuolization phenotype, no accumulation of APs is observed after 72 hr of STF treatment and the number of LAMP-1 positive structures is maintained (**Fig. 2D, 2E**). Confirming LC3-turnover assays, these results show that STF-62247 blocks later stages of autophagy and leads to a reduction in the number of endo-lysosomes in VHL-mutated RCC4 cells.

**Figure 2.**
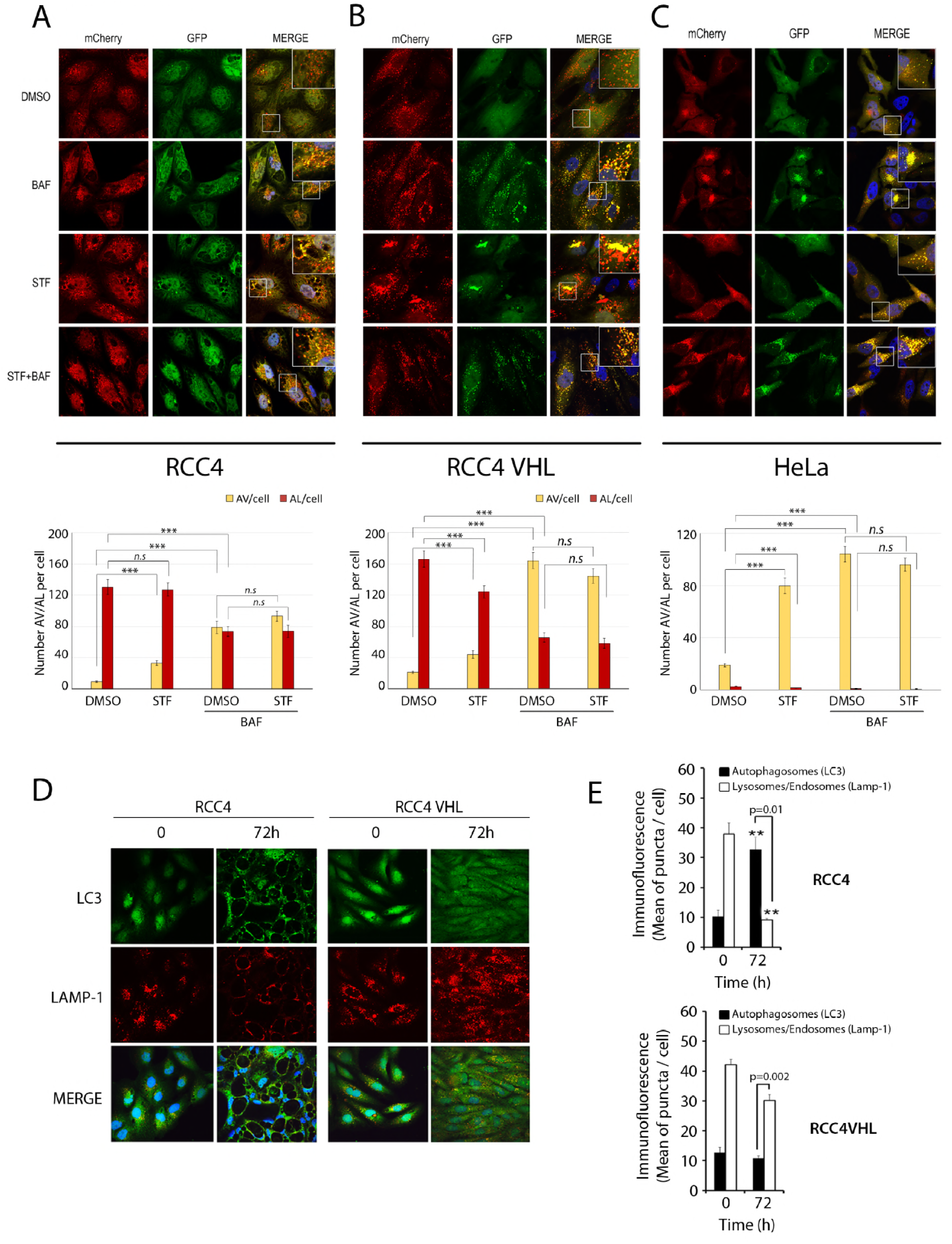
Long-term STF-62247 treatment decreases the number of endo-lysosomal vesicles in VHL-mutated RCC4 cells. Confocal microscopy of cells stably expressing mCherry-GFP-LC3 with quantification of numbers of autophagic vesicles (AV) or autolysosomes (AL) in response to 4 hr of STF and BAF treatment alone or combined in **(A)** RCC4, **(B)** RCC4 VHL and **(C)** HeLa. **(D)** Immunofluorescence of endogenous LC3 and LAMP1 after 72 hr of STF in RCC4 and RCC4 VHL. **(E)** Puncta quantification of (D) LC3 and LAMP-1. Mean and SEM were calculated from at least three independent experiment. Statistical analysis compared untreated cells with treated cells (*p<0.05, **p<0.01, ***p<0.001) using Student’s T-test.

### Upstream signaling and canonical formation of autophagosomes are not affected by STF-62247

Small molecules and diverse synthetic compounds have already been reported to be able to simultaneously block later stages of autophagy and induce AP formation.(24–28) For this reason, initial steps of AP biogenesis as well as upstream signaling cascades leading to autophagy initiation have been explored in order to fully study STF-62247’s effects on this pathway. The class III phosphatidylinositol 3-kinase (VPS34) is responsible for the synthesis of phosphatidylinositol-3-phosphate (PI(3)P) at the AP precursor membranes and local increases in PI(3)P recruits specific autophagic effectors such as WD-repeat protein interacting with phosphoinositides 1 (WIPI-1 /Atg18).(23, 29–31) WIPI-1 puncta formation was visualized and quantified in order to explore STF’s effects on early phagophore formation (**Fig. 3A, 3B**). As expected, control starvation conditions (EBSS) increased the number of WIPI-1 puncta, indicative of autophagy initiation in both VHL-mutated and -proficient cell models (**Fig 3A, 3B**). STF treatment alone or in combination with starvation conditions however, did not increase WIPI-1 puncta formation in either cell models when compared to DMSO or EBSS controls, respectively (**Fig. 3A,3B**). Downstream of WIPI-1 recruitment, the conjugation of ATG5-ATG12 was measured by immunoblot assays (**Fig. 3C**). Measuring free-ATG12 and conjugated ATG5-ATG12 is an informative technique to assess novel AP formation. Thus, conjugated ATG5-ATG12 was quantified at an early and a late time point (8 hr and 48 hr STF treatment) in VHL-defective RCC4 cells and VHL-proficient cell lines (**Fig. 3C**). Coinciding with WIPI-1 puncta count, no significant increase in ATG5-ATG12 conjugation was measured in RCC4, RCC4VHL or in HeLa cells at an early time point (**Fig. 3C**). In fact, the 48 hr-time point shows a decrease in ATG5-ATG12 conjugation in RCC4 VHL cells while no significant changes are reported in the VHL-defective RCC4 cells and in HeLa (**Fig. 3C**). These results indicate that STF-62247 does not affect early steps of canonical autophagosome formation.

**Figure 3.**
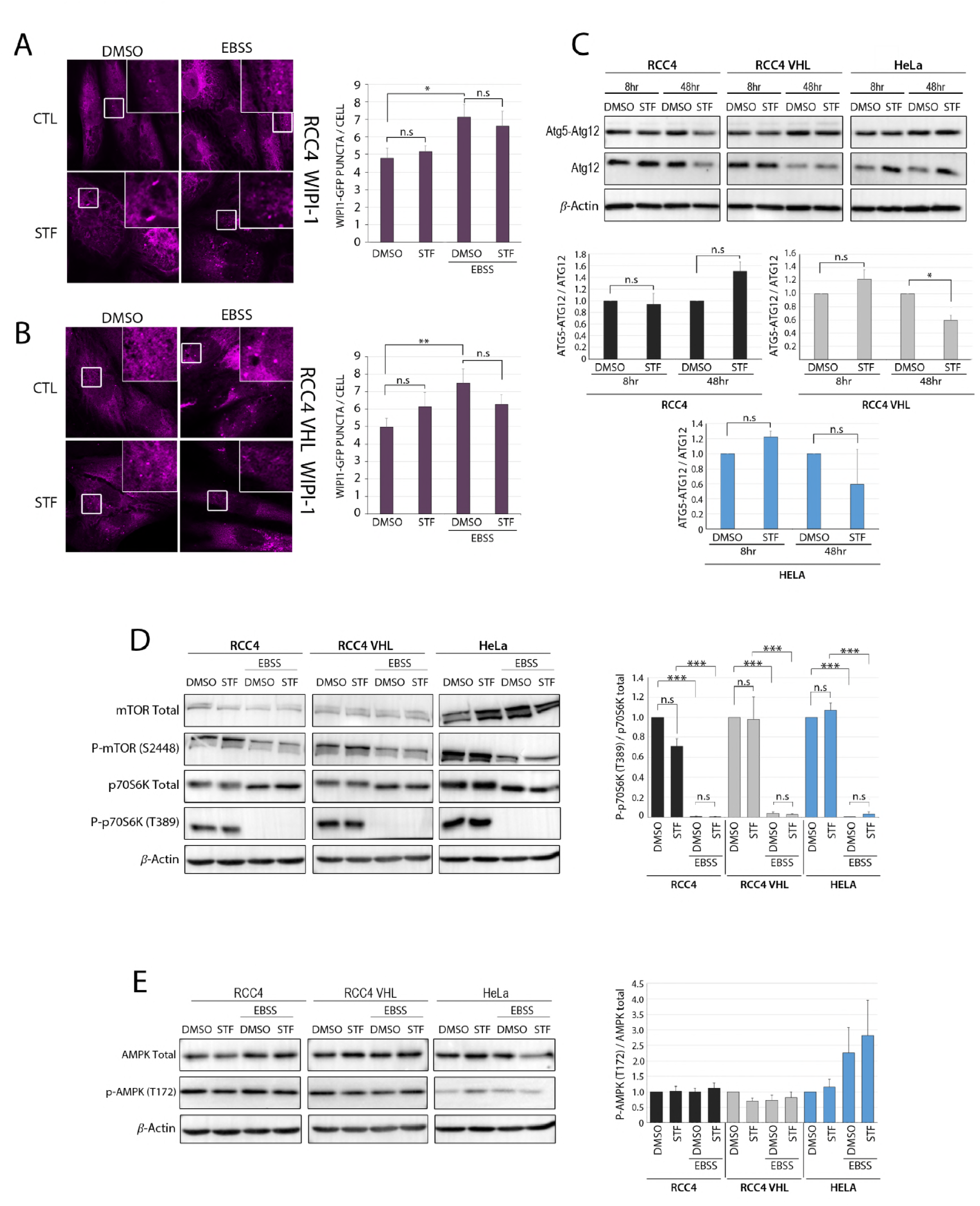
Upstream signaling and canonical formation of autophagosomes are not affected by STF-62247. Stable cells expressing GFP-WIPI-1 in response to 4 hr starvation conditions (EBSS) alone or combined with STF in **(A)** RCC4 and **(B)** RCC4 VHL. Number of puncta per cell was quantified for each condition. **(C)** Immunoblot analysis of levels of endogenous free ATG12 and ATG5-ATG12 complex after 8 hr or 48 hr of STF treatment in RCC4, RCC4 VHL and HeLa with corresponding quantification. **(D)**Immunoblot analysis of mTOR, P-mTOR(S2448) and of its downstream effectors p70S6K and P-p70S6K (T389) after 4hr of STF and starvation (EBSS) treatments alone, or combined in RCC4, RCC4VHL and HeLa cells with corresponding quantifications. **(E)** Immunoblot of total AMPK and p-AMPK (T172) in RCC4, RCC4 VHL and Hela, treated as seen in (D) and quantified. Mean and SEM were calculated from at least three independent experiment. Statistical analysis compared untreated cells with treated cells (*p<0.05, **p<0.01, ***p<0.001) using Student’s T-test.

A major suppressive regulator of canonical autophagy is the mammalian target of rapamycin (mTOR) complex 1 (mTORC1), activated at the lysosomal surface.(32, 33) Our previous study investigating STF-62247 by quantitative proteomics had highlighted a difference in mTOR activation between VHL-proficient and -deficient cell models in response to prolonged STF treatment times. Indeed, we highlighted the ability of VHL-proficient cells to restore mTOR activity following prolonged exposition to STF-62247 compared to VHL-mutated RCC4 cells.(34) To characterize earlier effects of STF on this signaling, immunoblot analyses were performed to measure phosphorylation states of mTOR and of its downstream effector RPS6KB1/p70S6 kinase in response to a 4 hr STF treatment (**Fig. 3D**). Control starvation conditions (EBSS) not only reduced the phosphorylation of P-mTOR (S2448), but completely inhibited the phosphorylation levels of P-p70S6K (T389) by mTOR (**Fig. 3D**). However, at variance with previously published prolonged STF-treatment times, 4 hr STF treatment alone or in combination with starvation conditions did not alter the levels of total mTOR and p70S6K or of their phosphorylation levels (**Fig. 3D**).

Next, total and phosphorylated states of AMP-activated protein kinase (AMPK), a multimeric serine/threonine protein kinase that acts as a fine-tuned sensor of energetic levels in the cell was measured in response to STF treatment. Phosphorylation of the conserved threonine residue (p-AMPK T172) is a prerequisite for its activity.(35) Consequently, activated AMPK inhibits mTOR function and acts as an inducer of autophagy via activation of the ULK1-mATG13-FIP200 complex. No significant alterations of total AMPK or p-AMPK T172 were observed in response to STF alone or when combined with starvation conditions in either VHL-mutated RCC4 cells or in VHL-proficient RCC4 VHL and Hela. These results indicate that while STF-62247 blocks later stages of autophagy, this small compound does not simultaneously affect upstream signaling or the canonical formation of autophagosomes.

### CRISPR/Cas9 knock-outs of essential ATG genes indicate a partial role for autophagy in STF-62247 signaling

Stable CRISPR/Cas9 knockout (KO) cell models of essential autophagy related genes (ATG) were established in VHL-mutated and -proficient cell lines to inhibit early steps of autophagy initiation and to elucidate the importance of this pathway in STF’s signaling. KOs of BECN-1/Beclin-1 and ATG5 were chosen to suppress autophagy as they play essential roles in the nucleation and maturation steps, respectively. (**Fig. 4A-4F**) XTT viability assays were performed in order to study the effects of these KOs on the sensitive VHL-mutated RCC4 cells in response to increasing concentrations of STF (**Fig 4A, 4B**). Impeding autophagy initiation in VHL-mutated RCC4 cells by crispr.Beclin-1 or crispr.ATG5 both rescued their viability and rendered them up to 56% and 46% more resistant at the highest STF concentration, respectively. Although an increase in viability is observed in VHL-defective RCC4 cells, this rescue is not found in VHL-proficient cell models, in fact it remains unchanged (**Fig. 4C-4F**). Moreover, the vacuolization phenotype was visualized in order to assess concomitant rescue. Intriguingly, only a partial rescue in phenotype was observed in VHL-mutated Crispr.Beclin-1 and Crispr.ATG5 cells, whilst no such rescue was observed in RCC4 VHL and HeLa (**Fig.4A-4B and 4C-4F**). Taken together, these results indicate that while autophagy plays a role in STF-62247 signaling in VHL-mutated RCC4 cells, the persisting phenotype alludes to contributions from additional intimately linked pathways.

**Figure 4.**
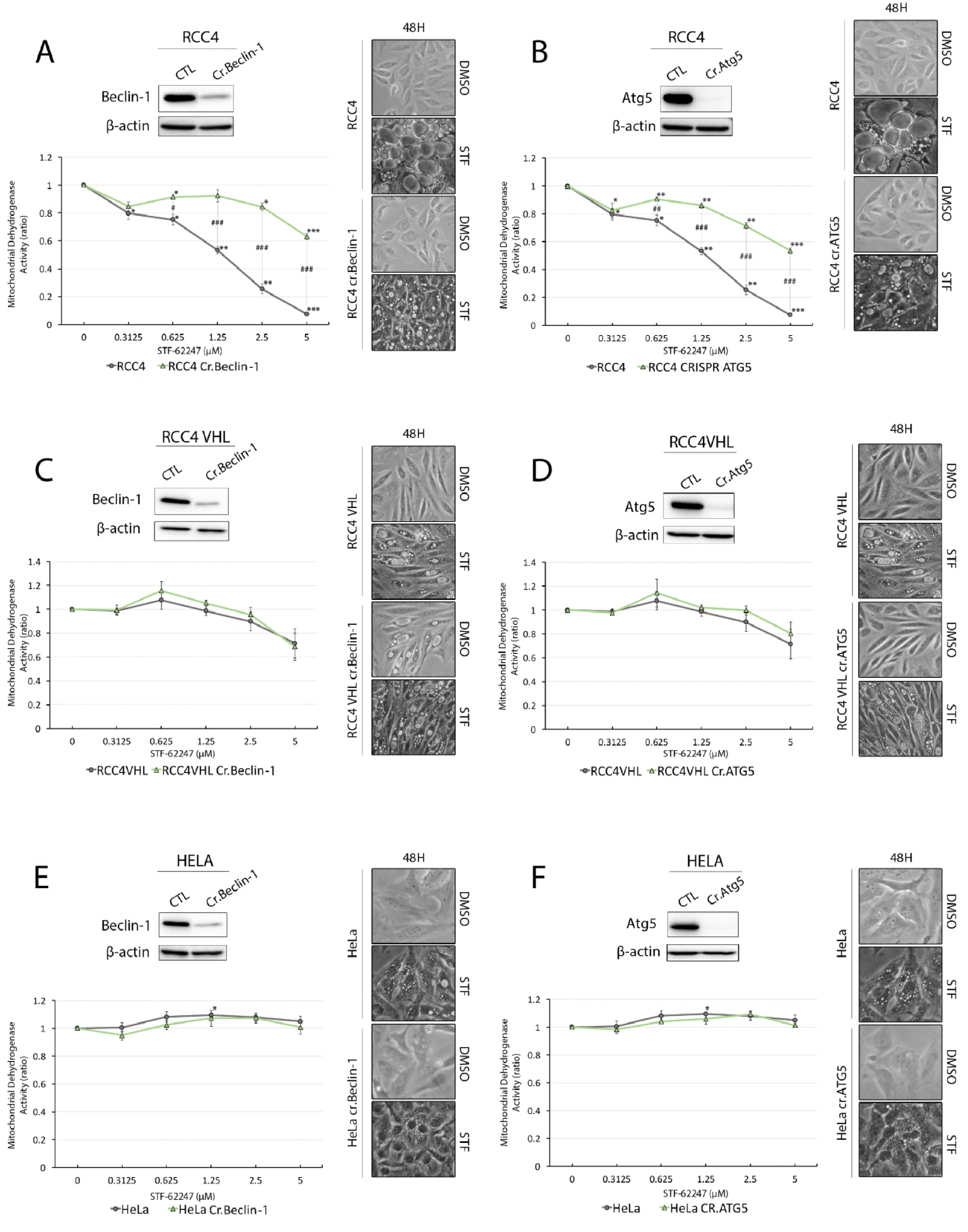
CRISPR/Cas9 knock-outs of essential ATG genes indicate a limited role for autophagy in STF-62247 signaling. CRISPR/Cas9 knock-out of essential autophagy-related genes; Beclin-1 and ATG5 with knock-out validation by immunoblot, XTT viability assay of CRISPR models in response to 48 hr STF treatment (0-5*μ*M) and corresponding inverted-light microscopy images in **(A-B)** RCC4, **(C-D)** RCC4 VHL and **(E-F)** Mean and SEM were calculated from at least three independent experiment. Statistical analysis compared untreated cells with treated cells (*p<0.05, **p<0.01, ***p<0.001) and the difference between the two cell lines (#p<0.05, ##p<0.01, ###p<0.001) at each concentration using Student’s T-test.

### STF-62247 localizes at lysosomes and induces swelling of endocytic compartments

The auto-fluorescent property of STF was exploited to visualize its cellular localization. Co-localization studies by live cell imaging of STF and tagged RFP-Bacmam transduced cells was accomplished to visualize endo-lysosomal and endoplasmic reticulum (ER) compartments (**Fig. 5A, 5B**). More specifically, CellLight lysosomes-RFP (RFP-Lamp-1) and ER-RFP (RFP-calreticulin and -KDEL sequence) were utilized to transduce VHL-mutated RCC4 and RCC4 VHL cells. The fluorescence of STF appears as defined green fluorescent puncta in both RCC4 and RCC4 VHL cells (**Fig. 5A, 5B**). Following treatment, STF almost entirely co-localizes with LAMP-1 positive compartments in both VHL-mutated and -proficient cell models (**Fig. 5A-5B**). Furthermore, the absence of STF fluorescent co-localization with ER compartments indicates that it does not bind randomly to a multitude of membrane-bound markers but is in fact specific to LAMP-1 positive structures (*i.e*. late endosomes and lysosomes) (**Fig 5A, 5B**).

**Figure 5.**
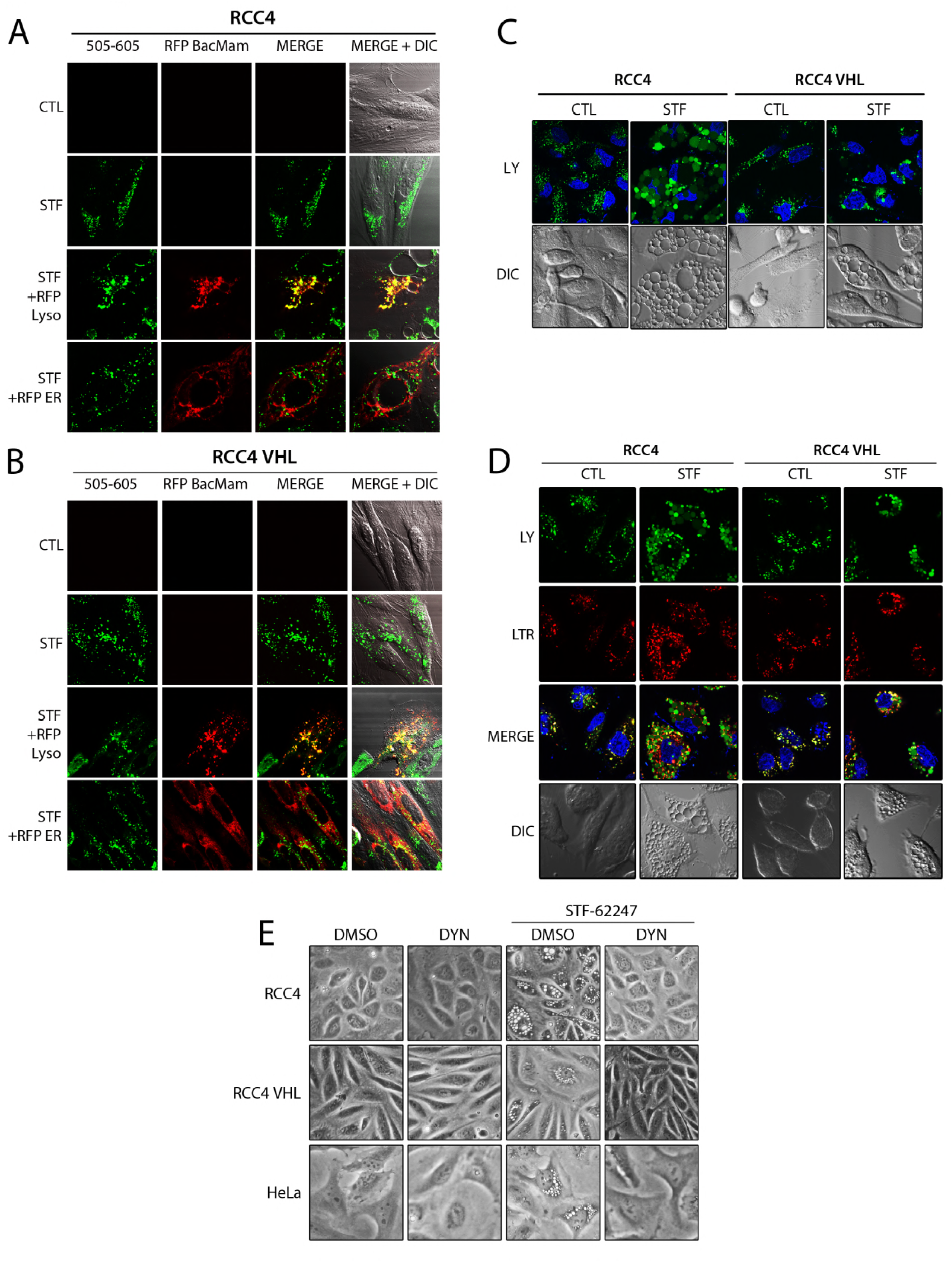
STF-62247 localizes at lysosomes and induces swelling of endocytic compartments. **(A-B)** RCC4 and Live-cell imaging of STF’s autofluorescence and tagged RFP-Bacmam transduced RCC4 and RCC4 VHL cells [CellLight lysosomes-RFP (RFP-Lamp-1) and ER-RFP (calreticulin and KDEL sequence] in response to 48 hr STF treatment. **(C)** Live-cell imaging of RCC4 and RCC4 VHL cells after 16 hr of Lucifer yellow internalization, 6 hr of STF treatment **(D)** and 30 min staining of Lysotracker. **(E)** Inverted-light microscopy images of RCC4, RCC4 VHL and Hela cells in response to 4 hr of Dynasore and STF treatments either alone or combined.

Lucifer Yellow (LY) is a well-known impermeable pH-insensitive marker of fluid phase endocytosis *i.e* pinocytosis/macropinocytosis; an endocytic mechanism for the non-specific bulk uptake and internalisation of extracellular fluid.(36–42) Considering that LY can only enter the cell by being internalized, this dye was an ideal tool to highlight the endocytic route, leading to late endosomes and lysosomes. Thus, in order to confirm the endo-lysosomal nature of these enlarged vacuoles, STF-treated cells were stained with LY (**Fig. 5C, 5D**). In RCC4 and RCC4 VHL DMSO controls, LY dye enters cells and forms fluorescent puncta-like structures (**Fig. 5C**). In STF-treated cells however, LY resides exclusively inside enlarged vacuoles confirming their endo-lysosomal origin (**Fig. 5C**). Lysotracker red (LTR) is a widely used acidotropic dye for the visualization of acidic organelles.(43) Both VHL-mutated and -proficient RCC models were treated with STF and imaged by live-cell microscopy after co-staining with LY and LTR red (**Fig. 5D**). The acidotropic LTR accumulates and decorates the limiting membranes of these LY-filled structures indicating possible deregulated fusion events between acidic compartments (**Fig. 5D**). The inability of LTR to stain these swollen organelles would lead us to believe that they might not be acidic in nature and in fact be earlier endocytic compartments. However, in similar cases of endo-lysosomal disruption and enlargement, others have also documented the inability of LTR red to mark the lumen of these swollen late endosome and lysosomal acidic compartments.(44–46)

Although blocking autophagy initiation through KOs of essential ATG genes significantly reduced RCC4 cells’ sensitivity to STF, they were unable to rescue the vacuolization phenotype (**Fig. 4A-4F**). For these reason, we hypothesized that STF might target endocytic pathways and have repercussions on autophagy termination. And so, a widely-used inhibitor of endocytic pathways was used (*i.e* Dynasore) in combination with STF and visualized by inverted-light microscopy (**Fig. 5E**). By visualizing VHL-mutated RCC4 cells as well as VHL-proficient RCC4 VHL and HeLa cell models, no changes in the cells’ appearance were documented in response to DYN treatment alone (**Fig. 5E**). As shown, treatment with STF immediately caused the swelling of endo-lysosomal compartments in all three cell models (**Fig. 5E**). However, when a simultaneous DYN treatment was added to STF, vacuolization was completely prevented and did not occur, resulting in a complete rescue of STF’s effect (**Fig. 5E**). Altogether, these results show that STF localizes at LAMP-1 positive structures (late-endosomes and lysosomes) and affects endocytosis through enlargement of endo-lysosomal structures.

### STF-62247 possesses lysosomotropic properties and affects lysosomal integrity and cathepsin D trafficking

The localization of STF at lysosomal structures led us to hypothesize that this compound had lysosomotropic characteristics and the vacuolization phenotype could in fact originate from the swelling of acidic endo-lysosomal structures. Similar to LTR, acridine orange (AO) is another acidotropic dye (AO) that was used to visualize the acidity of these translucid vacuoles. The mechanism of action of this weak base metachromatic dye is better characterized than that of LTR and is known for its capacity to stain acidic organelles such as late endosomes and lysosomes. AO molecules become protonated and trapped and their accumulation inside these acidic endo-lysosomal structures leads to a shift in excitation from green to red (530nm to 620nm) thus, permitting the differentiation of acidic organelles (red-orange granules) from other cellular compartments (diffuse green).(53–55) Following a short-term STF treatment, AO staining revealed vivid red fluorescence as a result of its rapid accumulation inside the growing vacuoles present in VHL-mutated RCC4 cells (**Fig. 6A**). No striking differences in the numbers of red granules were observed in STF-treated VHL-proficient cells compared to the DMSO control; concurrent with the modest vacuolization phenotype of these cells (**Fig. 6A**). Furthermore, BAF treatment was utilized as a specificity control for AO staining as it neutralizes endo-lysosomal pH via inhibition of the v-ATPase proton pump. As expected, BAF treatment alone eliminated the levels of red granules in both RCC4 and RCC4 VHL cell models (**Fig. 6A**). Moreover, the addition of BAF to STF treatment resulted in a complete loss of red fluorescence and seemed to rescue the endo-lysosomal swelling phenotype, indicating that STF could no longer have its effect when lysosomal pH was neutralized (**Fig. 6A**). To differentiate a true rescue in the phenotype from a possible inability of these vacuoles to become acidic, inverted light microscope images were taken in both VHL-mutated RCC4 cells and RCC4 VHL (**Fig. 6B**). The combination of BAF plus STF treatment completely inhibited the formation of STF-induced vacuoles resulting in a full phenotype rescue highlighting both the importance of the V-ATPase pump in STF’s signaling and the compounds lysosomotropic properties (**Fig. 6B**). AO staining was also performed at a later time point to see if the vacuoles retained their capacity to trap AO molecules (**Fig. 6C, 6D**). Even at 72 hr of STF treatment, these extremely swollen compartments retained their acidity as AO molecules were still accumulating within them, getting protonated and emitting red fluorescence (**Fig. 6C**). RCC4 VHL cells however, showed no differences in red granule staining in response to 72 hr of STF treatment (**Fig. 6D**).

**Figure 6.**
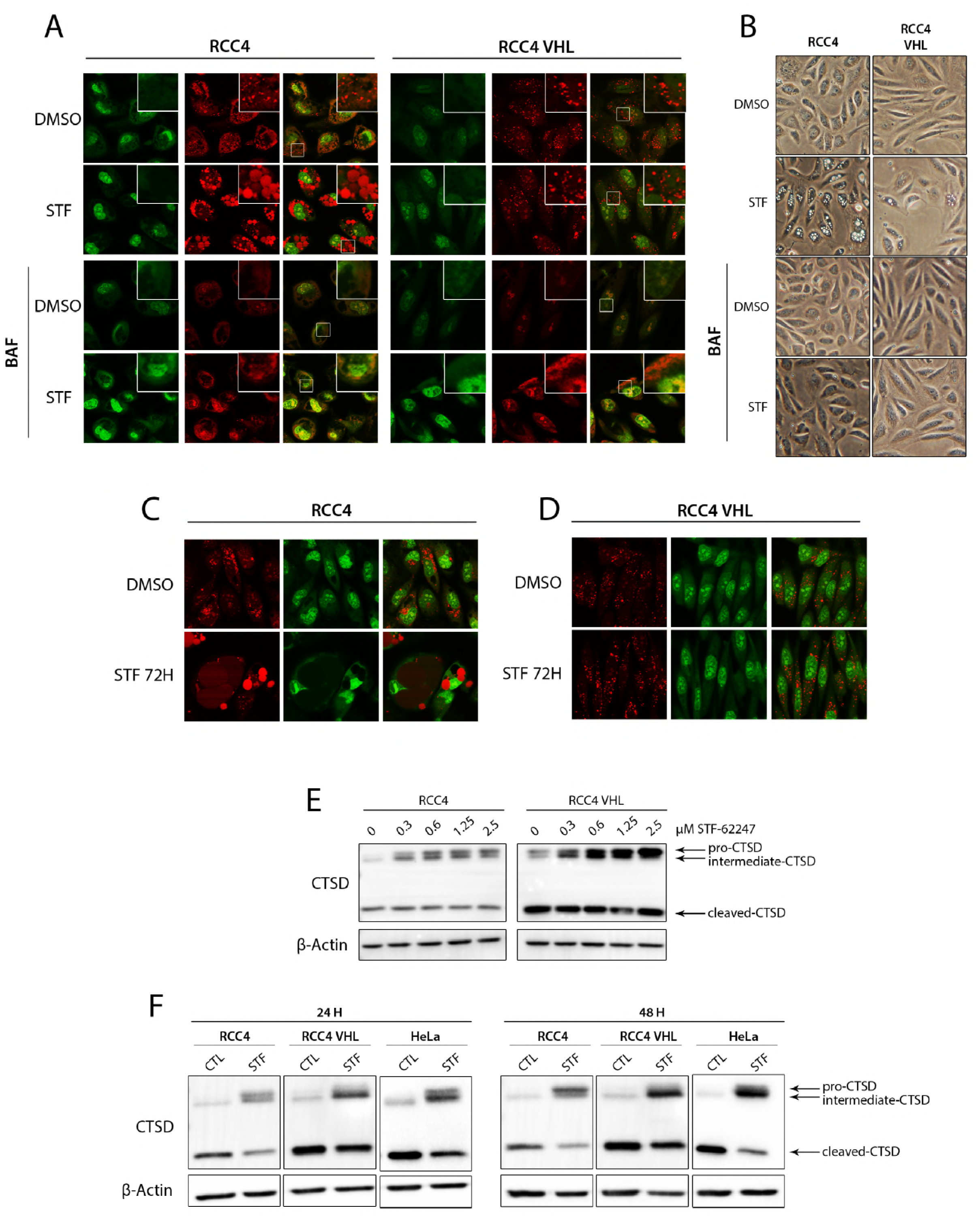
STF-62247 possesses lysosomotropic properties and affects lysosomal integrity and cathepsin D trafficking. **(A)** Live-cell images of RCC4 and RCC4 VHL treated 4 hr with STF and BAF alone or combined followed by 10 minutes of acridine orange staining. **(B)** Inverted-light microscopy images of RCC4 and RCC4 VHL models after 4 hr STF and BAF treatments alone or combined. **(C)** Live-cell images of RCC4 and **(D)** RCC4 VHL treated 72 hr and followed by 10 minutes of acridine orange staining. **(E)** Immunoblot assay of cathepsin D (CTSD) levels in RCC4 and RCC4 VHL models in response to 4 hr of increasing STF concentration (0-2.5*μ*M). **(F)** immunoblot analysis of CTSD levels in RCC4, RCC4 VHL and HeLa models after 24 hr and 48 hr of STF treatment.

The observed enlargement of endo-lysosomal compartments should theoretically disrupt normal cellular vesicular trafficking. This hypothesis was tested by following the maturation of a well-known lysosomal aspartyl protease Cathepsin D (CTSD) which has a step-wise maturation process; synthesized in the ER, transported to Golgi stacks, then to endosomes and becoming completely active in the lysosomal lumen (**Fig. 6E, 6F**).(56–58) Disruption of this highly regulated process would result in accumulation of immature CTSD which would incidentally affect lysosomal degradative capacity and explain the observed block in later stages of autophagy. Immunoblot assays of CTSD (pro-CTSD (golgi); intermediate CTSD (endosome) and cleaved/active CTSD (lysosomes)) in response to increasing concentrations of STF were performed (**Fig. 6E**). Accumulation of pro- and intermediate-CTSD was observed in a concentration-dependent manner in both RCC4 and RCC4 VHL cell models (**Fig. 6E**). Next, longer STF treatment-times were tested in all three cell line models (**Fig. 6F**). In response to 24 hr and 48 hr of STF treatment, levels of active cleaved-CTSD were strikingly decreased with an accompanying accumulation of pro- and intermediate-CTSD in all three cell line models, highlighting the effect of STF on vesicular trafficking (**Fig. 6F**). Taken together, these results show that swollen endo-lysosomal compartments retain their acidity in response to STF, even at 72 hr of STF treatment. Interestingly, the complete phenotype rescue observed with the combinatory treatment of BAF plus STF confirms lysosomotropic characteristics of STF. Finally, the accumulation of immature cathepsin D shows that STF affects lysosomal integrity and vesicular trafficking concomitant with the measured block in later stages of autophagy.

## DISCUSSION

The present work is the first to identify clear mechanistic insights into STF-62247’s effects on the autophagic flux and on its lysosomotropic properties. By utilizing several autophagic assays exploring both later stages of autophagy as well as important upstream signaling pathways leading to autophagy initiation, we show that STF-62247 is a clear blocker of later stages of autophagy through endocytic and vesicular trafficking deregulation and does not affect initiation of canonical AP formation.

Our results build on and enhance the comprehension of the original study that aimed at selectively targeting VHL-defective RCCs.(10). Originally identified as an autophagy inducer, we re-classify STF-62247 as a potent blocker of autophagy through disruption of lysosomal physiology. Initially, identification of STF-62247 was accomplished by high-throughput screening of 64 000 synthetic compounds in ccRCC models (RCC4 and its subclone counterpart expressing wild-type VHL; RCC4 VHL).(10, 20, 21, 59) With this in mind, cell models for this study were chosen very carefully. The identical cell models were chosen for a more thorough and accurate characterization of STF-62247’s effects on autophagy and its related signaling pathways. Moreover, HeLa cells were chosen as a control cell model for two reasons; firstly, because autophagy has been abundantly studied in this cell line and secondly, because it harbours endogenous wild-type VHL, differentiating it from the VHL over-expressing subclone RCC4 VHL.

By exploiting the natural auto-fluorescence of this small compound, we were able to pinpoint its cellular endo-lysosomal localization which leads to extreme enlargement of these structures in the VHL-mutated model. Unfortunately, even by localizing STF to LAMP-1 positive structures, we have yet to identify its protein target. This insight would be greatly beneficial in understanding the exact cause of the endo-lysosomal swelling phenotype, something we are actively pursuing. Although autophagy blockage by STF-62247 is independent of VHL-status, it would seem that the acclimatization to this small compound (*i.e*. capacity of VHL-positive cells to surmount lysosomal swelling) is in fact dependent on this tumor suppressor. These results strongly suggest a possible unidentified role for VHL in the maintenance of lysosomal integrity. Although this was not the objective of the present work, we highlighted the ability of VHL-containing cells to surmount the STF-induced enlargement of late endosomes and lysosomes and renew their numbers (LAMP-1 positive structures) at a prolonged STF treatment time; something VHL-mutated cells were unable to do. In fact, mTOR re-activation has been shown to be an essential step for autophagic lysosomal reformation, a fascinating mechanism of lysosomal recycling and reformation where tubules and vesicles emanate from autolysosomes and mature into newly functional lysosomes. (60–62) During this process and after prolonged stress conditions (or starvation), the autolysosomal degradation of macromolecules triggers the reactivation of mTOR in order to replenish the number of lysosomes. While short-term STF treatment does not seem to affect mTOR phosphorylation, we have previously shown that prolonged treatment of STF decreases the phosphorylation levels and activity of mTOR in VHL-deficient cells, concomitant with an inability of these cells to renew LAMP-1 positive structures. Moreover, in VHL-proficient cells, mTOR phosphorylation levels are returned to normal as compared to the DMSO control after prolonged STF treatment times.(34) These intriguing results merit further investigations as it is a possibility that the capability of VHL-proficient cells to surmount the endolysosomal swelling induced by STF-62247 could be linked to their ability to form new lysosomes. Elucidating these insights would help explain the toxicity differences observed between VHL-mutated and -functioning cells.

Autophagy has a multifaceted role in tumor evasion from immune surveillance and targeting its later stages shows great potential in ccRCCs.(63) VHL-defective tumours have an impressively high basal autophagic flux in response to stabilization of hypoxia inducible factors and blocking late stages of autophagy through disruption of vesicle trafficking and lysosomal integrity could be a promising strategy to re-sensitize resistant tumours to already available targeted therapies.(63–67) This study re-classifies STF-62247 as a potent blocker of autophagy through vesicular trafficking and lysosomal defects and we highlight the possibility of exploiting this small compound in autophagic and lysosomal research that could undeniably lead to a better fine-tuning of specific mechanisms of autophagic suppression for therapeutic purposes.

## METHODS/MATERIALS

### Cell culture and treatments

Parental ccRCC cell line (**RCC4)** and its subclone counterpart stably transfected with an expression vector encoding VHL (**RCC4 VHL)** were a gift from Dr. Amato Giaccia (Stanford University). Authentication of parental ccRCC cell lines was performed by short tandem repeat (STR) DNA profile at Genetica DNA Laboratories (Burlington, NC). Human cervical carcinoma cells (**HeLa**) were purchased ATCC (#*CCL-2*). All three cell lines were cultured in Dulbecco’s Modified Eagle Medium (DMEM) high glucose (GE healthcare, *#SH30081.01*) supplemented with 10% fetal bovine serum (Thermo Fisher, *#10082147*) 2mM L-glutamine (Fisher Scientific, *#SH30034.01*) and 1mM sodium pyruvate (GE healthcare, *#SH30239.01*). All cell lines tested negative for mycoplasma contamination.

Treatments were as follows; **STF-62247** (Cayman Chemical, *#13084*), 1.25*μ*M for 4 hr, 24 hr, 48 hr and 72 hr; **Bafilomycin A1** (Cayman Chemical, *#11038*), 600nM (RCC4, RCC4 VHL) and 400nM (HeLa) for 4 hr; **Dynasore** (Sigma Aldrich, *#D7693*), 200 *μ*M (RCC4, RCC4 VHL) and 100*μ*M (HeLa) for 4 hr. Starvation conditions (Earl’s Balanced Salt Solution (EBSS) (Thermo Fisher, *#24010043*), 4 hr at 37°C under 5% CO_2_.

### Plasmids, stable cell line constructions and CRISPR/Cas9 models

GFP-LC3 and mCherry-GFP-LC3 were amplified from pBABE-puro mCherry-EGFP-LC3B gifted from Jayanta Debnath (Addgene, *#22418*) using the primer pairs *5′-*CACCATGGT GAGCAAGGGCGA-3′ (forward); *5′-*TTACACTGACAATTTCATCCCGAA-3′ (reverse) and inserted in pLenti CMV Blast DEST (706–1), a gift from Eric Campeau & Paul Kaufman (Addgene, *#17451*).(68, 69) A lipidation defective LC3 mutant was generated by site-directed mutagenesis using the primer pairs *5′-*CACCATGGTGAGCAAGGGCGAGGAG-3′ (forward); 5′-TTACACTGACAATTTCATCGCGAACGTCTCCT-3′ (reverse) to convert the conserved C-terminal glycine 120 residue into an alanine (G→A) and was inserted in pLenti CMV/TO Puro DEST (670-1), a gift from Eric Campeau & Paul Kaufman (Addgene, # 17293). Crispr/Cas9 sgRNA sequences for Beclin-1 and Atg5 are the following; 5′-CACCATTCCATTCCACGGGAACAC-3′ (forward); 5-AAACGTGTTCCCGTGGAATGGAATC-3′ (reverse) and 5′-CACCTCAGAAGCTGTTTCGTCCTG-3′ (forward); 5′-AAACCA GGACGAAACAGCTTCTGAC-3′ (reverse), respectively. These sequences were inserted in lentiCRISPRv2 plasmid, a gift from Feng Zhang (Addgene; *#52961*) as previously described.(70, 71) The retroviral plasmid pMXs-IP GFP-WIPI-1 was a gift from Noboru Mizushima (Addgene; *#38272*).

### XTT viability assay

XTT viability assays were performed as previously described.(34) Briefly, five thousand cells were seeded per well in 96-well plates. Cells were treated with serial dilutions of STF-62247 after which plates were incubated at 37°C under 5% CO_2_ for 48 hr or 96 hr. Following the incubation period, XTT solution (comprising 0.3mg/mL of XTT powder (Sigma Aldrich, *#X4626*), DMEM high glucose without phenol red (Wisent Bio, *#319-051-CL*), 20% FBS and 2.65*μ*g/mL phenazine methosulfate (PMS) (Sigma Aldrich, *#P9625*)) was added to each well and plates were incubated at 37°C for 1 hr. Absorbance was then read at 450nm on a Spectramax Plus spectrophotometer (Molecular Devices, Sunnyvale, CA).

### Inverted-light microscopy imaging

Images of untreated and treated cells (vacuolization) were taken with a CoolPix990 Nikon camera (Nikon, Japan) mounted on a Nikon TMS-F inverted microscope (Nikon, Japan).

### Western Blot analysis

Western Blot were performed as previously described.(23, 34) Briefly, cells were lysed in M-PER buffer and quantified using Pierce BCA protein assay Kit (Thermo Scientific, *#23225*). Protein samples were boiled for 5-7 minutes at 95°C, separated on sodium dodecyl sulfate polyacrylamide gels (SDS-PAGE) and transferred onto polyvinylidene difluoride (PVDF) membranes. Chemiluminescence detection was performed using the ECL Prime Western Blotting detection reagent kit (GE Healthcare, *#RPN2232*) on a Chemi-Doc XRS+ imager (BIO-RAD Inc, Mississauga, ON, Canada). Primary antibodies used include β-Actin (*sc-47778*), Cathepsin D (*sc-6486*), p62/SQSTM1 (*sc-28359*) and Lamp-1 (immunofluorescence only; *sc-* 20011) from Santa Cruz Biotechnologies; LC3 (*#3868*), ATG12 (#*4180*), ATG5 (*#12994*), Beclin-1 (*#3495*), mTOR (*#2983*), phospho-mTOR(S2448) (*#5536*), p70S6K (*#2708*), phospho-p70S6K (T389) (*#9234*), AMPK (*#5832*), and phospho-AMPK (T172) (*#2535*) from Cell Signaling Technologies.

### Immunofluorescence

Cells were grown on coverslips at 30% confluency and were fixed in 3.7% formaldehyde (Sigma Aldrich, *#F8775*) and then permeabilized with 0.25% Triton x-100 in PBS. FBS was used for blocking, primary and secondary antibodies.(23) All images were taken on an Olympus Fluoview FV1000 confocal microscope (Olympus, Center Valley, Pennsylvania, USA) with a 60X oil-immersion lens.

### Lucifer Yellow, Lysotracker and Acridine Orange staining

Lucifer Yellow. 15 000 cells were seeded in Lab-Tek II 8-well chambered coverglass (Thermo Fisher, *#155409*) and loaded with 1 mg/ml of Lucifer Yellow (Biotium, *#80015*) for 16 hr.(39, 40) Cells were washed 3X with PBS and incubated with fresh media or with media containing 1.25 µM of STF-62247 for 6 hr. After 5.5 hr of incubation, 75 nM of Lysotracker Red (Molecular Probes, #*L7528)* and/or 2 µg/ml Hoechst (Molecular Probes, *#H1399*) were added to the media for 30 minutes. Finally, cells were washed 3X with PBS and placed in Phenol Red free DMEM media for live-cell imaging. A minimum of three images per condition were obtained for three independent experiments using an Olympus Fluoview FV1000 confocal microscope (Olympus, Center Valley, Pennsylvania, USA).

Acridine Orange (Sigma Aldrich, *#A6014*) staining was optimized from previously published studies.(17, 54, 72) Briefly, cells were treated with 1.25µM of STF-62247 for 4 hr or 72 hr after which they were loaded with a solution of 10ug/mL of AO for 10 minutes. Cells were washed 3X with PBS and incubated with pre-warmed Phenol Red free DMEM media and imaged using an Olympus Fluoview FV1000 confocal microscope (Olympus, Center Valley, Pennsylvania, USA).

### STF-62247 auto-fluorescence and localization assay

10 000 cells were seeded in in Lab-Tek II 8-well chambered coverglass. Cells were treated with 1.25µM STF-62247 for 16 hr. 6µL of Cell Light Lysosomes-RFP, BacMam 2.0 (Thermo Fisher, #C10504) and ER-RFP, BacMam 2.0 (Thermo Fisher, #*10591*) were added to STF-treated wells overnight (12 hr).(73) Cells were washed 3X with PBS and a fresh solution of 5µM STF-62247 in pre-warmed Phenol Red free DMEM media was added on the cells and immediately imaged using an Olympus Fluoview FV1000 confocal microscope (Olympus, Center Valley, Pennsylvania, USA).

### Quantification and Statistical Analyses

Statistical data are presented as mean ±SEM. The statistical significance is determined using two-tailed Student t test from a minimum of 3 independent experiments. Statistical analysis compared untreated cells with treated cells (*p<0.05, **p<0.01, ***p<0.001) as well as differences between cell lines (#p<0.05, ##p<0.01, ###p<0.001).

## ACKNOWLEDGMENTS

This work was supported by the grants from the Canadian Institutes of Health Research (CIHR) and the New Brunswick Innovation Foundation (NBIF). Sandra Turcotte is also supported by a research chair from the Canadian Cancer Society New-Brunswick.

## AUTHOR CONTRIBUTIONS

NB conceived the project and designed the experiments under the supervision of ST. NB performed all experiments in figure 1, 2 and 3, figure 5A-B and figure 6 (including live-cell confocal and immunofluorescence microscopy images acquisition but excluding puncta quantification). NB wrote the manuscript. DC was responsible for the generation of all plasmids, stable cell line constructions and CRISPR/Cas9 models. DC was responsible for all immunofluorescence puncta quantification (figure.1F-1G, figure.2A-C, figure.3A-B) and performed all experiments in figure 4 as well as figure 5B. KC performed the experiments of figure 5C-5D. ST performed the experiments in figure 2D and 2E and edited the manuscript with NB.

## List of abbreviations

AMPK: AMP-activated protein kinase
AO: Acridine Orange
AP: Autophagosome
ATG: Autophagy-related gene
BAF: Bafilomyin A1
ccRCC: Clear cell renal cell carcinoma
CTSD: Cathepsin D
KO: Knock-out
LTR: Lysotracker Red
LY: Lucifer Yellow
mTOR: Mammalian target of Rapamycin
STF: STF-62247
V-ATPase: Vacuolar H+-ATPase pump
VHL: von Hippel-Lindau

## REFERENCES

1. Kovacs G, Akhtar M, Beckwith BJ, Bugert P, Cooper CS, Delahunt B, et al. The Heidelberg classification of renal cell tumours. The Journal of pathology. 1997;183(2):131–3. doi: 10.1002/(SICI)1096-9896(199710)183:2<131::AID-PATH931>3.0.CO;2-G. PubMed PMID: 9390023.

2. Parikh M, Lara PN, Jr. Modern Systemic Therapy for Metastatic Renal Cell Carcinoma of the Clear Cell Type. Annual review of medicine. 2017. doi: 10.1146/annurev-med-041916-124132. PubMed PMID: 29144835.

3. Levy DA, Slaton JW, Swanson DA, Dinney CP. Stage specific guidelines for surveillance after radical nephrectomy for local renal cell carcinoma. The Journal of urology. 1998;159(4):1163–7. PubMed PMID: 9507823.

4. Maxwell PH, Wiesener MS, Chang GW, Clifford SC, Vaux EC, Cockman ME, et al. The tumour suppressor protein VHL targets hypoxia-inducible factors for oxygen-dependent proteolysis. Nature. 1999;399(6733):271–5. doi: 10.1038/20459. PubMed PMID: 10353251.

5. Motzer RJ, Agarwal N, Beard C, Bolger GB, Boston B, Carducci MA, et al. NCCN clinical practice guidelines in oncology: kidney cancer. J Natl Compr Canc Netw. 2009;7(6):618–30. PubMed PMID: 19555584.

6. Escudier B, Porta C, Schmidinger M, Algaba F, Patard JJ, Khoo V, et al. Renal cell carcinoma: ESMO Clinical Practice Guidelines for diagnosis, treatment and follow-up. Annals of oncology: official journal of the European Society for Medical Oncology / ESMO. 2014;25 Suppl 3:iii49–56. doi: 10.1093/annonc/mdu259. PubMed PMID: 25210086.

7. Vitale MG, Carteni G. Clinical management of metastatic kidney cancer: the role of new molecular drugs. Future Oncol. 2016;12(1):83–93. doi: 10.2217/fon.15.283. PubMed PMID: 26617188.

8. Xiaohong Ma SP, Quentin Mcafee and Ravi K. Amaravadi. Autophagy inhibition as a strategy for cancer therapy. In: Frederick R. Maxfield JMWaSL, editor. Lysosomes: Biology, Diseases, and Therapeutics. 1st ed: John Wiley & Sons, Inc.; 2016.

9. Maes H, Rubio N, Garg AD, Agostinis P. Autophagy: shaping the tumor microenvironment and therapeutic response. Trends in molecular medicine. 2013;19(7):428–46. doi: 10.1016/j.molmed.2013.04.005. PubMed PMID: 23714574.

10. Turcotte S, Chan DA, Sutphin PD, Hay MP, Denny WA, Giaccia AJ. A molecule targeting VHL-deficient renal cell carcinoma that induces autophagy. Cancer cell. 2008;14(1):90–102. doi: 10.1016/j.ccr.2008.06.004. PubMed PMID: 18598947; PubMed Central PMCID: PMC2819422.

11. Colecchia D, Stasi M, Leonardi M, Manganelli F, Nolano M, Veneziani BM, et al. Alterations of autophagy in the peripheral neuropathy Charcot-Marie-Tooth type 2B. Autophagy. 2017:0. doi: 10.1080/15548627.2017.1388475. PubMed PMID: 29130394.

12. Conte A, Paladino S, Bianco G, Fasano D, Gerlini R, Tornincasa M, et al. High mobility group A1 protein modulates autophagy in cancer cells. Cell death and differentiation. 2017;24(11):1948–62. doi: 10.1038/cdd.2017.117. PubMed PMID: 28777374; PubMed Central PMCID: PMCPMC5635219.

13. Wan W, You Z, Xu Y, Zhou L, Guan Z, Peng C, et al. mTORC1 Phosphorylates Acetyltransferase p300 to Regulate Autophagy and Lipogenesis. Molecular cell. 2017;68(2):323–35 e6. doi: 10.1016/j.molcel.2017.09.020. PubMed PMID: 29033323.

14. Su H, Yang F, Wang Q, Shen Q, Huang J, Peng C, et al. VPS34 Acetylation Controls Its Lipid Kinase Activity and the Initiation of Canonical and Non-canonical Autophagy. Molecular cell. 2017;67(6):907–21 e7. doi: 10.1016/j.molcel.2017.07.024. PubMed PMID: 28844862.

15. Xie Z, Xie Y, Xu Y, Zhou H, Xu W, Dong Q. Bafilomycin A1 inhibits autophagy and induces apoptosis in MG63 osteosarcoma cells. Mol Med Rep. 2014;10(2):1103–7. doi: 10.3892/mmr.2014.2281. PubMed PMID: 24890793.

16. Yamamoto A, Tagawa Y, Yoshimori T, Moriyama Y, Masaki R, Tashiro Y. Bafilomycin A1 prevents maturation of autophagic vacuoles by inhibiting fusion between autophagosomes and lysosomes in rat hepatoma cell line, H-4-II-E cells. Cell structure and function. 1998;23(1):33–42. PubMed PMID: 9639028.

17. Yoshimori T, Yamamoto A, Moriyama Y, Futai M, Tashiro Y. Bafilomycin A1, a specific inhibitor of vacuolar-type H(+)-ATPase, inhibits acidification and protein degradation in lysosomes of cultured cells. The Journal of biological chemistry. 1991;266(26):17707–12. PubMed PMID: 1832676.

18. Klionsky DJ, Abdelmohsen K, Abe A, Abedin MJ, Abeliovich H, Acevedo Arozena A, et al. Guidelines for the use and interpretation of assays for monitoring autophagy (3rd edition). Autophagy. 2016;12(1):1–222. doi: 10.1080/15548627.2015.1100356. PubMed PMID: 26799652; PubMed Central PMCID: PMCPMC4835977.

19. Szeto J, Kaniuk NA, Canadien V, Nisman R, Mizushima N, Yoshimori T, et al. ALIS are stress-induced protein storage compartments for substrates of the proteasome and autophagy. Autophagy. 2006;2(3):189–99. PubMed PMID: 16874109.

20. Hay MP, Turcotte S, Flanagan JU, Bonnet M, Chan DA, Sutphin PD, et al. 4-Pyridylanilinothiazoles that selectively target von Hippel-Lindau deficient renal cell carcinoma cells by inducing autophagic cell death. Journal of medicinal chemistry. 2010;53(2):787–97. doi: 10.1021/jm901457w. PubMed PMID: 19994864; PubMed Central PMCID: PMC2838715.

21. Turcotte S, Sutphin PD, Giaccia AJ. Targeted therapy for the loss of von Hippel-Lindau in renal cell carcinoma: a novel molecule that induces autophagic cell death. Autophagy. 2008;4(7):944–6. PubMed PMID: 18769110; PubMed Central PMCID: PMC2803726.

22. Shi B, Huang QQ, Birkett R, Doyle R, Dorfleutner A, Stehlik C, et al. SNAPIN is critical for lysosomal acidification and autophagosome maturation in macrophages. Autophagy. 2017;13(2):285–301. doi: 10.1080/15548627.2016.1261238. PubMed PMID: 27929705; PubMed Central PMCID: PMCPMC5324844.

23. Vicinanza M, Korolchuk VI, Ashkenazi A, Puri C, Menzies FM, Clarke JH, et al. PI(5)P regulates autophagosome biogenesis. Molecular cell. 2015;57(2):219–34. doi: 10.1016/j.molcel.2014.12.007. PubMed PMID: 25578879; PubMed Central PMCID: PMCPMC4306530.

24. Rez G, Csak J, Fellinger E, Laszlo L, Kovacs AL, Oliva O, et al. Time course of vinblastine-induced autophagocytosis and changes in the endoplasmic reticulum in murine pancreatic acinar cells: a morphometric and biochemical study. European journal of cell biology. 1996;71(4):341–50. PubMed PMID: 8980904.

25. Ostenfeld MS, Hoyer-Hansen M, Bastholm L, Fehrenbacher N, Olsen OD, Groth-Pedersen L, et al. Anti-cancer agent siramesine is a lysosomotropic detergent that induces cytoprotective autophagosome accumulation. Autophagy. 2008;4(4):487–99. PubMed PMID: 18305408.

26. Cheong H, Lindsten T, Wu J, Lu C, Thompson CB. Ammonia-induced autophagy is independent of ULK1/ULK2 kinases. Proceedings of the National Academy of Sciences of the United States of America. 2011;108(27):11121–6. doi: 10.1073/pnas.1107969108. PubMed PMID: 21690395; PubMed Central PMCID: PMCPMC3131371.

27. Eng CH, Yu K, Lucas J, White E, Abraham RT. Ammonia derived from glutaminolysis is a diffusible regulator of autophagy. Science signaling. 2010;3(119):ra31. doi: 10.1126/scisignal.2000911. PubMed PMID: 20424262.

28. Wu YT, Tan HL, Shui G, Bauvy C, Huang Q, Wenk MR, et al. Dual role of 3-methyladenine in modulation of autophagy via different temporal patterns of inhibition on class I and III phosphoinositide 3-kinase. The Journal of biological chemistry. 2010;285(14):10850–61. doi: 10.1074/jbc.M109.080796. PubMed PMID: 20123989; PubMed Central PMCID: PMCPMC2856291.

29. Simonsen A, Tooze SA. Coordination of membrane events during autophagy by multiple class III PI3-kinase complexes. The Journal of cell biology. 2009;186(6):773–82. doi: 10.1083/jcb.200907014. PubMed PMID: 19797076; PubMed Central PMCID: PMCPMC2753151.

30. Jaber N, Dou Z, Chen JS, Catanzaro J, Jiang YP, Ballou LM, et al. Class III PI3K Vps34 plays an essential role in autophagy and in heart and liver function. Proceedings of the National Academy of Sciences of the United States of America. 2012;109(6):2003–8. doi: 10.1073/pnas.1112848109. PubMed PMID: 22308354; PubMed Central PMCID: PMCPMC3277541.

31. Jean S, Kiger AA. Classes of phosphoinositide 3-kinases at a glance. Journal of cell science. 2014;127(Pt 5):923–8. doi: 10.1242/jcs.093773. PubMed PMID: 24587488; PubMed Central PMCID: PMCPMC3937771.

32. Sancak Y, Bar-Peled L, Zoncu R, Markhard AL, Nada S, Sabatini DM. Ragulator-Rag complex targets mTORC1 to the lysosomal surface and is necessary for its activation by amino acids. Cell. 2010;141(2):290–303. doi: 10.1016/j.cell.2010.02.024. PubMed PMID: 20381137; PubMed Central PMCID: PMCPMC3024592.

33. Sancak Y, Peterson TR, Shaul YD, Lindquist RA, Thoreen CC, Bar-Peled L, et al. The Rag GTPases bind raptor and mediate amino acid signaling to mTORC1. Science. 2008;320(5882):1496–501. doi: 10.1126/science.1157535. PubMed PMID: 18497260; PubMed Central PMCID: PMCPMC2475333.

34. Bouhamdani N, Joy A, Barnett D, Cormier K, Leger D, Chute IC, et al. Quantitative proteomics to study a small molecule targeting the loss of von Hippel-Lindau in renal cell carcinomas. International journal of cancer Journal international du cancer. 2017;141(4):778–90. doi: 10.1002/ijc.30774. PubMed PMID: 28486780.

35. Alers S, Loffler AS, Wesselborg S, Stork B. Role of AMPK-mTOR-Ulk1/2 in the regulation of autophagy: cross talk, shortcuts, and feedbacks. Molecular and cellular biology. 2012;32(1):2–11. doi: 10.1128/MCB.06159-11. PubMed PMID: 22025673; PubMed Central PMCID: PMCPMC3255710.

36. Stewart WW. Lucifer dyes--highly fluorescent dyes for biological tracing. Nature. 1981;292(5818):17–21. PubMed PMID: 6168915.

37. Swanson JA, Yirinec BD, Silverstein SC. Phorbol esters and horseradish peroxidase stimulate pinocytosis and redirect the flow of pinocytosed fluid in macrophages. The Journal of cell biology. 1985;100(3):851–9. PubMed PMID: 3972898; PubMed Central PMCID: PMCPMC2113515.

38. Riezman H. Endocytosis in yeast: several of the yeast secretory mutants are defective in endocytosis. Cell. 1985;40(4):1001–9. PubMed PMID: 3886157.

39. Basrai MA, Naider F, Becker JM. Internalization of lucifer yellow in Candida albicans by fluid phase endocytosis. J Gen Microbiol. 1990;136(6):1059–65. doi: 10.1099/00221287-136-6-1059. PubMed PMID: 2200841.

40. Wiederkehr A, Meier KD, Riezman H. Identification and characterization of Saccharomyces cerevisiae mutants defective in fluid-phase endocytosis. Yeast. 2001;18(8):759–73. doi: 10.1002/yea.726. PubMed PMID: 11378903.

41. Baluska F, Samaj J, Hlavacka A, Kendrick-Jones J, Volkmann D. Actin-dependent fluid-phase endocytosis in inner cortex cells of maize root apices. J Exp Bot. 2004;55(396):463–73. doi: 10.1093/jxb/erh042. PubMed PMID: 14739268.

42. Buckley CM, King JS. Drinking problems: mechanisms of macropinosome formation and maturation. The FEBS journal. 2017;284(22):3778–90. doi: 10.1111/febs.14115. PubMed PMID: 28544479.

43. Johnson DE, Ostrowski P, Jaumouille V, Grinstein S. The position of lysosomes within the cell determines their luminal pH. The Journal of cell biology. 2016;212(6):677–92. doi: 10.1083/jcb.201507112. PubMed PMID: 26975849; PubMed Central PMCID: PMCPMC4792074.

44. Jefferies HB, Cooke FT, Jat P, Boucheron C, Koizumi T, Hayakawa M, et al. A selective PIKfyve inhibitor blocks PtdIns(3,5)P(2) production and disrupts endomembrane transport and retroviral budding. EMBO Rep. 2008;9(2):164–70. doi: 10.1038/sj.embor.7401155. PubMed PMID: 18188180; PubMed Central PMCID: PMCPMC2246419.

45. Ho CY, Choy CH, Wattson CA, Johnson DE, Botelho RJ. The Fab1/PIKfyve phosphoinositide phosphate kinase is not necessary to maintain the pH of lysosomes and of the yeast vacuole. The Journal of biological chemistry. 2015;290(15):9919–28. doi: 10.1074/jbc.M114.613984. PubMed PMID: 25713145; PubMed Central PMCID: PMCPMC4392288.

46. Hazeki K, Uehara M, Nigorikawa K, Hazeki O. PIKfyve regulates the endosomal localization of CpG oligodeoxynucleotides to elicit TLR9-dependent cellular responses. PloS one. 2013;8(9):e73894. doi: 10.1371/journal.pone.0073894. PubMed PMID: 24040108; PubMed Central PMCID: PMCPMC3767827.

47. Girard E, Paul JL, Fournier N, Beaune P, Johannes L, Lamaze C, et al. The dynamin chemical inhibitor dynasore impairs cholesterol trafficking and sterol-sensitive genes transcription in human HeLa cells and macrophages. PloS one. 2011;6(12):e29042. doi: 10.1371/journal.pone.0029042. PubMed PMID: 22205993; PubMed Central PMCID: PMCPMC3242776.

48. Robinet P, Fradagrada A, Monier MN, Marchetti M, Cogny A, Moatti N, et al. Dynamin is involved in endolysosomal cholesterol delivery to the endoplasmic reticulum: role in cholesterol homeostasis. Traffic. 2006;7(7):811–23. doi: 10.1111/j.1600-0854.2006.00435.x. PubMed PMID: 16787396.

49. Grimmer S, Ying M, Walchli S, van Deurs B, Sandvig K. Golgi vesiculation induced by cholesterol occurs by a dynamin- and cPLA2-dependent mechanism. Traffic. 2005;6(2):144–56. doi: 10.1111/j.1600-0854.2005.00258.x. PubMed PMID: 15634214.

50. Weller SG, Capitani M, Cao H, Micaroni M, Luini A, Sallese M, et al. Src kinase regulates the integrity and function of the Golgi apparatus via activation of dynamin 2. Proceedings of the National Academy of Sciences of the United States of America. 2010;107(13):5863–8. doi: 10.1073/pnas.0915123107. PubMed PMID: 20231454; PubMed Central PMCID: PMCPMC2851890.

51. Preta G, Lotti V, Cronin JG, Sheldon IM. Protective role of the dynamin inhibitor Dynasore against the cholesterol-dependent cytolysin of Trueperella pyogenes. FASEB journal: official publication of the Federation of American Societies for Experimental Biology. 2015;29(4):1516–28. doi: 10.1096/fj.14-265207. PubMed PMID: 25550455; PubMed Central PMCID: PMCPMC4396600.

52. Preta G, Cronin JG, Sheldon IM. Dynasore - not just a dynamin inhibitor. Cell Commun Signal. 2015;13:24. doi: 10.1186/s12964-015-0102-1. PubMed PMID: 25889964; PubMed Central PMCID: PMCPMC4396812.

53. Lovelace MD, Cahill DM. A rapid cell counting method utilising acridine orange as a novel discriminating marker for both cultured astrocytes and microglia. Journal of neuroscience methods. 2007;165(2):223–9. doi: 10.1016/j.jneumeth.2007.06.009. PubMed PMID: 17662460.

54. Uchimoto T, Nohara H, Kamehara R, Iwamura M, Watanabe N, Kobayashi Y. Mechanism of apoptosis induced by a lysosomotropic agent, L-Leucyl-L-Leucine methyl ester. Apoptosis: an international journal on programmed cell death. 1999;4(5):357–62. PubMed PMID: 14634338.

55. Sohaebuddin SK, Tang L. A simple method to visualize and assess the integrity of lysosomal membrane in mammalian cells using a fluorescent dye. Methods Mol Biol. 2013;991:25–31. doi: 10.1007/978-1-62703-336-7_3. PubMed PMID: 23546655.

56. Gieselmann V, Hasilik A, von Figura K. Processing of human cathepsin D in lysosomes in vitro. The Journal of biological chemistry. 1985;260(5):3215–20. PubMed PMID: 3972822.

57. Laurent-Matha V, Derocq D, Prebois C, Katunuma N, Liaudet-Coopman E. Processing of human cathepsin D is independent of its catalytic function and auto-activation: involvement of cathepsins L and B. Journal of biochemistry. 2006;139(3):363–71. doi: 10.1093/jb/mvj037. PubMed PMID: 16567401; PubMed Central PMCID: PMCPMC2376303.

58. Hamer I, Jadot M. Endolysosomal transport of newly-synthesized cathepsin D in a sucrose model of lysosomal storage. Experimental cell research. 2005;309(2):284–95. doi: 10.1016/j.yexcr.2005.06.006. PubMed PMID: 16055118.

59. Reyjal J, Cormier K, Turcotte S. Autophagy and cell death to target cancer cells: exploiting synthetic lethality as cancer therapies. Advances in experimental medicine and biology. 2014;772:167–88. doi: 10.1007/978-1-4614-5915-6_8. PubMed PMID: 24272359.

60. Shen HM, Mizushima N. At the end of the autophagic road: an emerging understanding of lysosomal functions in autophagy. Trends Biochem Sci. 2014;39(2):61–71. doi: 10.1016/j.tibs.2013.12.001. PubMed PMID: 24369758.

61. Rong Y, McPhee CK, Deng S, Huang L, Chen L, Liu M, et al. Spinster is required for autophagic lysosome reformation and mTOR reactivation following starvation. Proc Natl Acad Sci U S A. 2011;108(19):7826–31. doi: 10.1073/pnas.1013800108. PubMed PMID: 21518918; PubMed Central PMCID: PMCPMC3093520.

62. Yu L, McPhee CK, Zheng L, Mardones GA, Rong Y, Peng J, et al. Termination of autophagy and reformation of lysosomes regulated by mTOR. Nature. 2010;465(7300):942–6. doi: 10.1038/nature09076. PubMed PMID: 20526321; PubMed Central PMCID: PMC2920749.

63. Janji B, Viry E, Moussay E, Paggetti J, Arakelian T, Mgrditchian T, et al. The multifaceted role of autophagy in tumor evasion from immune surveillance. Oncotarget. 2016;7(14):17591–607. doi: 10.18632/oncotarget.7540. PubMed PMID: 26910842; PubMed Central PMCID: PMCPMC4951235.

64. Messai Y, Noman MZ, Janji B, Hasmim M, Escudier B, Chouaib S. The autophagy sensor ITPR1 protects renal carcinoma cells from NK-mediated killing. Autophagy. 2015:0. doi: 10.1080/15548627.2015.1017194. PubMed PMID: 25714778.

65. Messai Y, Noman MZ, Hasmim M, Janji B, Tittarelli A, Boutet M, et al. ITPR1 protects renal cancer cells against natural killer cells by inducing autophagy. Cancer research. 2014;74(23):6820–32. doi: 10.1158/0008-5472.CAN-14-0303. PubMed PMID: 25297632.

66. Lotze MT, Maranchie J, Appleman L. Inhibiting autophagy: a novel approach for the treatment of renal cell carcinoma. Cancer J. 2013;19(4):341–7. doi: 10.1097/PPO.0b013e31829da0d6. PubMed PMID: 23867516.

67. Liang X, De Vera ME, Buchser WJ, Romo de Vivar Chavez A, Loughran P, Beer Stolz D, et al. Inhibiting systemic autophagy during interleukin 2 immunotherapy promotes long-term tumor regression. Cancer research. 2012;72(11):2791–801. doi: 10.1158/0008-5472.CAN-12-0320. PubMed PMID: 22472122; PubMed Central PMCID: PMCPMC3417121.

68. N’Diaye EN, Kajihara KK, Hsieh I, Morisaki H, Debnath J, Brown EJ. PLIC proteins or ubiquilins regulate autophagy-dependent cell survival during nutrient starvation. EMBO Rep. 2009;10(2):173–9. doi: 10.1038/embor.2008.238. PubMed PMID: 19148225; PubMed Central PMCID: PMCPMC2637314.

69. Campeau E, Ruhl VE, Rodier F, Smith CL, Rahmberg BL, Fuss JO, et al. A versatile viral system for expression and depletion of proteins in mammalian cells. PloS one. 2009;4(8):e6529. doi: 10.1371/journal.pone.0006529. PubMed PMID: 19657394; PubMed Central PMCID: PMCPMC2717805.

70. Ran FA, Hsu PD, Wright J, Agarwala V, Scott DA, Zhang F. Genome engineering using the CRISPR-Cas9 system. Nature protocols. 2013;8(11):2281–308. doi: 10.1038/nprot.2013.143. PubMed PMID: 24157548; PubMed Central PMCID: PMC3969860.

71. Sanjana NE, Shalem O, Zhang F. Improved vectors and genome-wide libraries for CRISPR screening. Nat Methods. 2014;11(8):783–4. doi: 10.1038/nmeth.3047. PubMed PMID: 25075903; PubMed Central PMCID: PMCPMC4486245.

72. Erdal H, Berndtsson M, Castro J, Brunk U, Shoshan MC, Linder S. Induction of lysosomal membrane permeabilization by compounds that activate p53-independent apoptosis. Proceedings of the National Academy of Sciences of the United States of America. 2005;102(1):192–7. doi: 10.1073/pnas.0408592102. PubMed PMID: 15618392; PubMed Central PMCID: PMCPMC544072.

73. Ames RS, Kost TA, Condreay JP. BacMam technology and its application to drug discovery. Expert opinion on drug discovery. 2007;2(12):1669–81. doi: 10.1517/17460441.2.12.1669. PubMed PMID: 23488908.

